# Interactions and information: Exploring task allocation in ant colonies using network analysis

**DOI:** 10.1101/2021.03.29.437501

**Authors:** Anshuman Swain, Sara D. Williams, Louisa J. Di Felice, Elizabeth A. Hobson

**Author notes:** contributed equally (Anshuman Swain, Sara D. Williams, Louisa J. Di Felice).

## Abstract

In animal societies, individuals may take on different roles to fulfil their own needs and the needs of their groups. Ant colonies display high levels of organisational complexity, with ants fulfilling different roles at different timescales (what is known as *task allocation*). Factors affecting task allocation can be at the individual level (e.g., physiology), or at the group level (e.g., the network of interactions). We focus on group level processes by exploring the relationship between interaction networks, task allocation and task switching using a previously published dataset (Mersch et al., 2013) tracking the behaviour of six *Camponotus fellah* colonies over 41 days. In our new analyses, our goal was to better explain the noisy process of task switching beyond simple age polyethism. First, we investigated the architecture of interaction networks using node (individual) level network measures and their relation to the individual’s task – foraging, cleaning or nursing – and whether or not the ant switched tasks. We then explored how noisy information propagation was among ants, as a function of the colony composition (how many ants carried out which tasks), through the information-theoretic metric of *Effective Information*. Our results show that interaction history was tied to task allocation: ants who switched to a task are more likely to have interacted with other ants carrying out that task. The degree to which interactions related to task allocation, as well as the noise in those interactions, depended on which groups of ants were interacting. Overall, we showed that colony cohesion was stable even as ant-level network measures varied more for ants when they switched functional groups; thus, ant colonies maintained a high level of information flow as determined by network analysis, and ant functional groups played different roles in maintaining colony cohesion through varied information flows.

**Highlights:**

- We analysed the interaction networks of six *Camponotus fellah* colonies
- We tested how centrality and information flow were tied to task switching
- Node-level network metrics and the information theoretic measure of *Effective Information* explained differences among functional groups
- Interactions were correlated with task switching, but the strength of the correlation differed across functional groups

## INTRODUCTION

In animal societies, individuals may carry out different tasks to fulfil their own needs and the needs of their group (Sumpter, 2006; Clutton-Brock, 2009; Jeanson & Weidenmuller, 2014). Larger and more complex societies can self-organise to fulfil tasks beyond basic sustenance and reproduction (Boomsma and Frank, 2006; Sumpter, 2010). Local exchange of information, between individuals of a group and between individuals and their environment, is key to self-organisation (Sumpter, 2006; Boomsma and Frank, 2006; Couzin, 2009; Cavagna et al., 2010; Swain and Fagan, 2019). Social insect colonies display high levels of organisational complexity (Lukas & Clutton-Brock, 2018), where individual tasks may include foraging, nest construction, and caring for the young (Gordon, 2002). The assignment of tasks, also referred to as *task allocation*, is the result of patterns of factors that vary across different scales (Gordon, 2015). These tasks can be fixed throughout each individual’s lifetime due to physiological reasons, for example when only a fertile subset of the population is responsible for reproduction, or when a subset is responsible for providing food (Sumpter, 2010; Clutton-Brock et al., 2001).

Task allocation can also result in individuals changing their main task over time. Task allocation in ants has been the subject of much previous work (Anderson and Shea, 2001; Gordon, 2015). Across ant species, studies have shown that, depending on the tasks and on the colony, ants may display varying degrees of task flexibility, from small colonies of totipotent ants to larger ones with a structured division of labour (Anderson and Shea, 2001). Factors affecting task changes can occur at the individual level or at the group level. Individual-level factors include physiology (Anderson and Shea, 2001), age (Tripet and Nonacs, 2004), corpulence (Robinson et al., 2009) and past experience (Ravary et al., 2007), whereas group-level factors involve colony size (Ravary et al., 2007) and interaction rates at the colony level (Gordon and Mehdiabadi, 1999). Studying individual-level factors associated with task change is often simpler than studying group-level ones. For example, individual-level changes can be easier to track because their rate of change often follows a consistent and predictable pattern, as in the case of ageing. Individual-level factors can also be directly quantified, e.g., by measuring age, corpulence, or physiological features, and traditional statistical approaches can be used to predict task changes.

An ant’s propensity to switch to a new task could also be linked to nature of its interactions and the topology of the group’s social interaction network structure. However, changes in task allocation affected by group-level factors are currently not well understood. Structural features or macro-level social properties of groups can affect micro-level individual actions if the social system is affected by feedbacks (Flack, 2017; Hobson et al., 2019). However, quantifying relevant macro-to-micro feedbacks can be challenging and can require large amounts of data. The development of automated tracking systems has made this level of data collection possible. While these systems have improved researchers’ ability to track detailed social behaviour (Robinson et al., 2009; Smith and Pinter-Wollman, 2021), assigning quantitative metrics to group dynamics is still a non-trivial task. In the case of interaction patterns, tracking physical interactions among individuals does not necessarily map onto the amount of meaningful (predictive) information exchanged with each interaction (Valentini et al., 2020). Although tracking technologies can tell us how many times individuals in a social group interact with one another, they cannot explain to what extent these interactions are tied to task allocation without considering the structure of these interactions and without including behavioural observations. Network methods and metrics allow us to explore the interaction structure.

In this paper, we leverage social network methods to gain new insight into task allocation changes in an existing dataset of ant interactions (published by Mersch et al. 2013). Mersch et al. studied task switching in *Camponotus fellah* by tracking and analysing the movements and interactions of individually-identified ants. Worker ants were categorised into three *functional groups* (nurse, cleaner or forager). Analyses showed that ants had more interactions with others in their same functional group. Communities defining the functional groups exhibited distinct behavioural signatures and were highly spatially divided. Nurses spent most of their time with the brood, while foragers spent time at the nest entrance and cleaners were located between the other two groups and the rubbish pile (Mersch et al. 2013). Mersch et al. also explored the questions of task switching cost, i.e., a time and energy investment associated with learning new tasks (Goldsby et al., 2012), and of age polytheism, i.e., the correlation between the age of an ant and which task they perform. The original study identified spatial fidelity as a key regulator of ant social organisation and interaction frequency (Mersch et al. 2013). They also found that task switches were present but uncommon and that when a shift in functional group occurred, ants showed a preferred direction of task transition, from nurses to cleaners to foragers, mostly based on age (Mersch et al., 2013). Task changes were thus hypothesised to be driven by age polyethism, but the patterns were fairly noisy.

In this new analysis we focus specifically on this noisy process of task switching and its predictability. A question not addressed in the previous study is whether the history of an ant’s interactions with others could be one of the elements explaining task switching. In other species, information flow patterns have been shown to affect task allocation and overall colony behaviour, such as in the case of midden workers in red harvester ants (*Pogonomyrmex barbatus*; Gordon and Mehdiabadi, 1999, Pinter-Wollman et al., 2018), tandem running recruitment (Franklin and Franks, 2012) and consensus-forming in rock ants (*Temnothorax albipennis*; Sasaki & Pratt, 2018). To test whether the history of interactions or information flow could explain the noise seen in task switching dynamics that was not explained by age polyethism alone, we evaluated several potential macro-scale predictors of task switching not addressed in the original paper.

First, we described the architecture of the interaction networks by focusing on information flow (which in our case refers to the possible information exchange due to interactions among ants). We tested whether the role played by individual ants in regulating information flow in the colony and the functional group that they belong to were correlated. To do this, we quantified three network measures that are tied to the architecture of information flows at the local level for ant-to-ant interactions: strength mode, betweenness centrality, and bridge betweenness centrality. We also quantified a network level measure, *Effective Information* (EI), for the whole colony. At the scale of ant-to-ant interactions, strength measures the quantity and frequency of interactions of an ant, and strength mode finds the value in the distribution of strengths most commonly observed across all the ants in the group. Betweenness centrality measures the number of shortest paths between pairs of ants that pass through it. Bridge betweenness centrality extends betweenness to measure the number of shortest paths that pass through a node and connect separate highly connected groups of nodes, or communities. While strength, betweenness centrality, and bridge betweenness centrality are common node-level measures in network science and have been applied to animal social networks in the past (Holme et al., 2002; Lusseau and Newman, 2004; Krause et al., 2009; Farine and Whitehead, 2015), Effective Information is a new information theoretic metric reflecting how noisy a mechanism connecting nodes (ants, in our case) is within a system. It is calculated as the difference between degeneracy and determinism of the network (Hoel, Albantakis, & Tononi, 2013; Klein and Hoel 2020). In interaction networks, Effective Information reflects the noisiness and predictability of the interactions among individuals (Hoel et al., 2020): a higher Effective Information means that a system is more deterministic, with information spreading in a more effective way throughout the network.

Second, we tested whether these four measures of information flow in the interaction network were related to task switching, to better understand the noise in task allocation not explained by age polyethism as determined by Mersch et al. (2013). We hypothesised that an ant’s previous interactions with other ants affect switching behaviour and tested whether interacting with a certain functional group increased the probability of an ant switching to that group. We found that the relationship between the structure of the interaction network and the different functional groups, as described by network measures at the node and the global level, could explain the varying correlations between interaction history and switching behaviour during task allocation. Our use of network metrics, including the Effective Information metric, allowed us to determine the relationship between interaction history, task allocation and information flow among functional groups in *Camponotus fellah* colonies.

## METHODS

### Data, network construction, and ant categorisation

The published Mersch et al. (2013) dataset contains summaries of interactions among a total of 985 individually-marked ants in six *Camponotus fellah* colonies. The authors collected interaction data for every pair of ants at a daily resolution over the 41-day monitoring period, and the published dataset contains data pooled at the number of interactions per dyad per day per colony. We matched this published dataset with the colony metadata to inform our analyses (Supplementary material 1).

Consistent with Mersch et al. (2013), we used the pairwise daily number of interactions to construct separate weighted, undirected, unipartite networks for each colony per day. Each ant in a colony was represented by an individual node. An edge between two nodes represents the interactions between those two ants on a given day. The edge weight is proportional to the number of pairwise interactions between them on that particular day. We used the available published dataset to recreate the 246 observed networks for the 6 colonies over 41 days used by Mersch et al. (2013) as well as the general pattern of task switching across the length of the experiments.

Mersch et al. (2013) assessed each ant’s functional group every 10 days to categorise them as a nurse, cleaner, or forager, representing their main task in the colony. They assigned functional groups based on what community an ant spent at least 70% of their time in, using the ‘infomap’ community detection algorithm paired with behavioural observations. They split the ants into the functional groups foragers (F), cleaners (C), nurses (N), queen (Q), and NA for ants who were counted as missing at a time point (e.g., if they were dead or had lost their tags).

Mersch et al. (2013) reported that their ants mostly did not change their task affiliation within the 10-day observation period between task assessment points. We used the same 10-day snapshot window in our analyses which resulted in three time points at which a switch in task to a new category could be detected. Based on Mersch et al’s (2013) observational data, when an ant switches functional groups, it switches tasks to that of the new group. For our analyses, we categorised each ant as “switched” or “consistent”, depending on whether they were categorised as part of a different functional group, or remained within the same functional group after each task assessment point in the original behavioural data. These labels were assigned for each 10-day observation period, meaning that an ant could be labelled as “consistent” in one time period because it did not change tasks from the previous period, and “switched” in the next if it then changed tasks, and thus functional groups during that next period. We utilised these labels and the functional groups set by Mersch et al. (2013), throughout our work.

Before performing new analyses, we first investigated whether we could replicate Mersch et al.’s (2013) results of age polyethism. We also tested whether we could recapitulate Mersch et al.’s results about task switching by determining the likelihood that an ant would stay in the same task throughout the experimental time versus performing two or three tasks.

### Quantifying individual network metrics for each ant

Our new analyses focused first at the individual scale within the networks. Node metrics and centralities define various types of influence that individual nodes exert on network connectivity and dynamics. For each network, we used R (v 3.6.2) and the packages *igraph* (Csardi and Nepusz, 2006) and *networktools* (Jones, 2020) to calculate three node-level, local metrics: (1) strength, (2) betweenness centrality, and (3) bridge betweenness. Since networks were constructed for each daily set of interaction observations for each colony, these metrics were calculated for each ant in every colony, every day. Differences in these metrics were then analysed as a function of functional groups at the colony level, and for just ants that switched or ants that remained consistent. First, we calculated each ant’s node strength as the sum of the weights of its edges.

Thus, in our context, it is a measure of not only how many interactions (edges) an ant (node) had with other ants, but also of how frequently those interactions occurred during a day. While degree is an index of potential communication activity (Freeman, 1979), strength improves upon this index by weighting degrees according to communication frequency, to better inform total interaction and information flow potential. To measure the structure of the distribution of this node level metric at the network level, we calculated the maxima of the density distribution of strength of all ants (or all within a functional group subset) in a given colony on a given day to find the *strength mode*. The mode was used instead of the mean because the strength distributions were skewed. The strength mode provides a summary of how these strengths are generally distributed across each network.

Second, we calculated each ant’s node betweenness. Also known as betweenness centrality, this measure is another way to assess the influence of a node for the connectivity of the network. For a given pair of nodes in a weighted network, there exists at least one path between them such that the sum of the link weights is minimized, thus forming a shortest path. The betweenness of a node is therefore defined as the number of shortest paths that pass through it. Freeman (1979) identified high betweenness centrality as a key indicator of whether a node occupies a central location in the network for information transmission. Individuals with high betweenness are often responsible for maintenance of communication, group coordination, and network stability (Lusseau and Newman, 2004; Farine and Whitehead, 2015). An ant with a high betweenness is an ant that is centrally located in the network, serving as a key connection for seemingly disparate ants.

Third, we measured the bridge betweenness for each ant in the network. Bridge betweenness extends the betweenness centrality metric to the level of communities and is defined as the number of times a node lies on the shortest path between two nodes from different communities. In network science, a community is defined as a group of nodes that have a higher likelihood of connecting to each other than to nodes from other communities. Ants with a high bridge betweenness serve as key connectors for different communities in the network, where communities mostly overlap with functional groups. This means that ants with high bridge betweenness would be more integral to network cohesion and information flow across groups, thus they may play an important role in driving switching dynamics.

To quantify each ant’s bridge betweenness, we needed to assign ants to network communities in both the observed networks but also in our 123,000 reference networks (see below). Assigning ants to network communities using the original network community detection algorithm Infomap (used in Mersch et al. 2013) was computationally prohibitive when applied to our many reference networks. Because of the computational demands of the bridge betweenness analysis, we used a Louvain community detection algorithm (Csardi and Nepusz, 2006) which saved computational time and memory (Emmons et al., 2016) compared to the Infomap algorithm. These new network community assignments were solely used for computing bridge betweenness and did not change the functional group assignments of the ants made by Mersch et al. (2013), which we use in all other cases in our analyses. To check that the Louvain algorithm assigned ants to network communities in ways consistent with the original community assignments from Infomap, we compared our community assignments to those found by Mersch et al. (2013): as we show below, our new assignments were similar enough to the original assignments that we could use our new method to assess bridge betweenness and the likelihood ants would be connected to others within different functional groups (see results, below). All other analyses involving functional group assignments of ants use the functional groups assigned in Mersch et al. (2013).

### Quantifying global network measures for each colony

We used *Effective Information* and its normalised measure, *Effectiveness*, to measure colony-level noisiness in the system, with respect to its underlying mechanisms (Hoel et al., 2020; Klein et al., 2022). Since we are considering the mechanism of communication and information flow among ants, Effective Information measures the level of predictability, or degeneracy, in ant-to-ant interactions. To calculate Effectiveness, we first found the sum of weights of all edges connected to each node in the interaction network, where edge weights correspond to the number of interactions between a pair of ants. An ant who had no interactions in a given network would have a weight of 0. We defined this weight as a vector *W*_*i*_ of the same length as the total number of nodes and referred to each element as *ω*_*ij*_, signifying the normalised value of edge weight between nodes *i* and *j*, such that for any index *i*, ∑_*j*_ ω_ij_ = 1. Here, each term *ω*_*ij*_ can be seen as the probability of moving from *i* to *j*, if a random walker is on the node *i*. Next, we characterised the uncertainty associated with each node *i*, calculated using Shannon’s entropy measure *H*(*W*_*i*_). As node *i* has more connections, and as the weights *ω*_*ij*_*s* of those connections to other nodes (*j*) become more equal, Shannon’s entropy (i.e., the uncertainty about where a random walker would go) increases. The average of this value across all the nodes in the network is < *H*(*W*_*i*_) >. When < *H*(*W*_*i*_) > is equal to 0, the network is deterministic (e.g., in the case of a line network or a ring lattice, both in directed and undirected cases, where information can only flow in one dimension). We then assessed the certainty of the network by calculating the term *H*(< *W*_*i*_ >), which is Shannon’s entropy of the average out-weights from all nodes. When this expression is equal to 0, the network is degenerate, with all edges leading to the same node. Finally, we calculated Effective Information using the following equation:

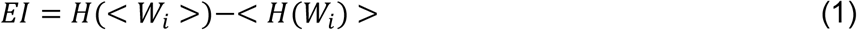

Here, the first term of the equation is determinism, and the second is degeneracy. Thus, the Effective Information for a line graph or a ring lattice, which are maximally deterministic and minimally degenerate, is the maximum. For the cases of a star network, which is both maximally deterministic and degenerate, and that of a complete graph, which is both minimally deterministic and degenerate, the value of Effective Information is zero. As the value of Effective Information can depend on the size of the network (Klein and Hoel, 2020), we calculated Effectiveness, the normalised Effective Information with respect to network size. Effective Information is normalized by *log*_*2*_*N*, which is the maximal possible value of the entropy, where *N* is the number of nodes in the network. For comparison, this is akin to the normalisation of Shannon diversity to Shannon equitability in ecological studies.

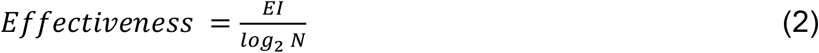

Effectiveness was calculated for each observed network (i.e., for each day, for every colony), using the R package *einet* (Byrum et al., 2020; Klein et al., 2022).

If node level properties were affected by which functional group an ant was in, then system level measures could be affected by the proportion of ants in each functional group, so we tested how group composition affected Effective Information. We used the functional group assignment from Mersch et al. (2013), then fit linear models to the Effectiveness for each observed network as a function of the proportion of each functional group in the network to determine significant relationships between Effectiveness and a colony’s functional group composition.

### Building reference models to test interaction patterns and task switching

To test how interaction patterns resulted in different network metrics and/or task switching patterns than expected, we constructed randomised networks that served as null models, or reference networks, for the daily interaction networks of the six colonies. Using randomised networks as reference networks is a common method for testing the effect of interaction structure and significance on various network properties and dynamics (Hobson et al. 2021; Farine 2017). Constructing a reference model allowed us to randomise some aspects of the interaction patterns while preserving other relevant structural features of the networks (Hobson et al. 2021). We used a degree-based randomisation (through the R package *VertexSort*; Abd-Rabbo, 2017) to generate our reference networks. This approach preserved the total number of interaction partners per any one ant on a given day but changed (1) who they interacted with and (2) how many times they interacted. This process distributed the total number of original interactions among the newly constructed edges of the randomised network.

This reference model approach allowed us to test how specific ant interaction patterns affected the node and network level properties, while preserving the distribution of connections in relation to functional groups of the ants (functional group assignment and the degree of individual ants remained unchanged). As an example, a nurse that had 20 interaction partners (degree = 20) would still have a degree of 20 in the reference network but would be interacting with 20 different ants with different frequencies, as the edge weights were also randomly assigned from the initial distribution for each reference network. This hypothetical reference model ant would then have a different total frequency of interactions while maintaining their original number of partners. For reproducibility, we created 500 seeded reference networks for each colony’s daily interaction network, for a total of 123,000 reference networks.

To test how observed network measures differed from those expected with the identity of interaction partners and the number of interactions randomised, we compared the observed node-level network measures to the distribution of those measures in our reference networks. We found the strength mode, mean betweenness, and mean bridge betweenness for every observed ant network (each colony, for each day) and for each of the reference networks. We also estimated the variance for each metric for every ant in a given colony on each day for the observed network and all the reference networks associated with the observed one (please note that in the case of strength, the variance is the standard variance in the strength distribution). The metrics were investigated separately for each functional group within the following subsets: overall (all ants), switching and consistent ants. Variance measured the individual variation of metrics among ants of one group in a colony. The distribution of variances, in conjunction with those of the central tendencies, helps us to explore the variation of these metrics across colonies and through time. Central tendencies (mode for strength, and mean for betweenness and bridge betweenness) and variance values were Z-transformed, separately for each metric and individual observed network with its respective reference networks, to facilitate comparison across observed networks which can vary in size (number of ants), allowing us to find each metric’s value for a given group, colony and day relative to its own reference models. The Z-transformation allowed us to combine values across colonies and days, and to visualize the means of those metrics across all samples (groups, days and colonies). We then calculated the 95% confidence intervals of the Z-transformed values for each functional group and ant subset (switching, consistent and overall ants) to determine differences in the network measures in relation to ant role and task switching. Since the distribution of variances indicates the variation of a network metric both across colonies and through time, the size of the 95% confidence intervals of the variances provides a proxy metric for stability of the network metrics, when interpreted in conjunction with the distribution of the central tendencies of those metrics.

To test if the frequency of interactions with different functional groups significantly affected an ant’s functional group membership and whether these interactions could explain how ants switched tasks, we compared the observed patterns of interactions in relation to switching behaviour to that of the reference models. If the functional group identity of ants affected how individuals contributed to information flow within a colony, then the number of times an ant switched to a new functional group should affect the flow of information. We tested how observed patterns differed from reference networks that preserved the number of interactions per ant but redistributed the interactions among every ant. We also tested whether the frequency of interactions with different functional groups significantly affected an ant’s final functional group. At each task assessment point, we quantified the frequency of interactions with each functional group before switching from its original functional group to the final one in both the observed dataset and in the ensemble of randomised reference networks. We compared the distribution of values computed from the observed networks against those given by the reference network distribution using a suite of chi-square (independence/homogeneity) comparisons separately for each possible type of task transition (including non-transitions) and each observed network, wherever the specific transition/non-transition occurred. To use the chi-square test, we assume that each interaction is independent as it involves transfer of new information between two individuals, even if it might be biased by more contact with certain individuals by choice. In addition, individual recognition is not particularly well established for ants, although they can recognize brood mates and their colony queen (see Esponda and Gordon, 2015; Sprenger and Menzel, 2020), lending more credibility to our assumption.

The significance results from each type of transition across all observed networks were then combined using Stouffer’s method (see Heard and Rubin-Delanchy, 2017) and significant differences at the alpha level of 0.05 were noted after accounting for multiple comparisons (across transitions) through the Benjamini-Hochberg (BH) correction. We combined all the cases where specific transitions were present, across all colonies and days. Importantly, we were comparing the total number of interactions in these tests pooled across all ants for a given subset, and not the number of ants that switched or remained consistent in their tasks. In all cases, the frequency of interactions with each type of functional group exceeded the minimum number required for using chi-square tests, even though in certain cases the number of ants who were interacting were less than five – their frequency of interactions exceed that number by a several orders of magnitude. This process allowed us to assess whether interactions within the previous observation period predicted functional group in the next observation period.

## RESULTS

### Replication of original results and visualisation of task switching

Figure 1 shows a new visualisation that summarizes the tasks of ants within all six colonies and how those tasks changed over time (Figure 1; for details, see Supplementary material 2, Table S1). To ground our analyses, we first replicated the main results from the original Mersch et al. (2013) paper. We were able to replicate the original results of age polyethism and recapitulated the distribution of the age of an ant that would switch tasks once, twice, or three times (results from Mersch et al., novel visualisation in Supplementary material 2, Figure S1A). We were also able to replicate Mersch et al.’s results about task switching by determining the likelihood that an ant would stay in the same task throughout the experimental time versus performing two or three tasks (Supplementary material 2, Figure S1B).

**Figure 1:**
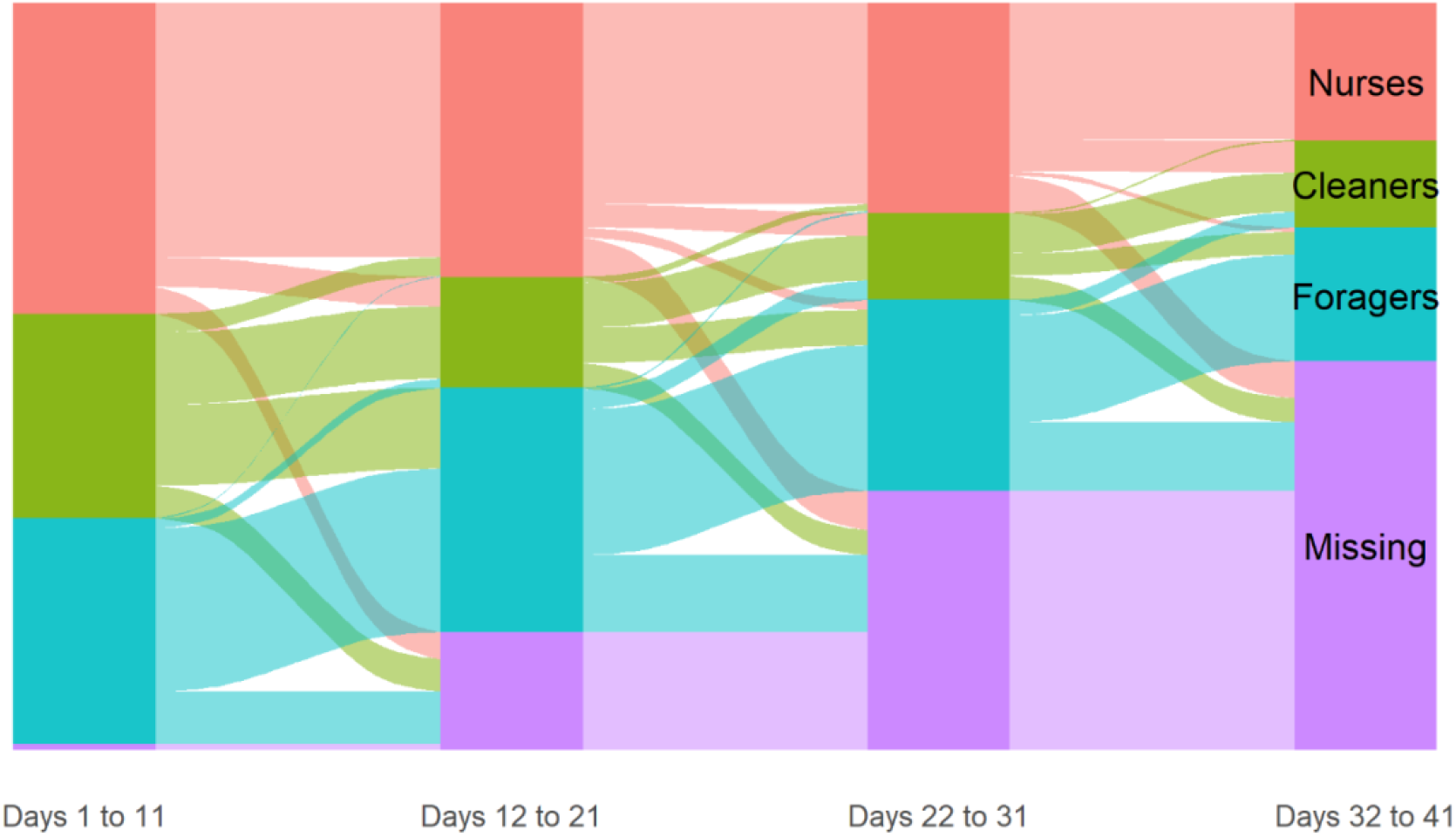
Dynamics of task allocation across the experimental time for all ants in all six colonies. The alluvial diagram shows the number of ants per functional group and number of ants staying in the same group or transitioning to a new functional group between time periods as proportional to box and flow sizes, respectively (functional groups as originally determined in Mersch et al. 2013 via infomap).

We also compared our interaction community assignments used in calculating bridge betweenness (via the Louvain community detection algorithm) with those obtained through the Infomap algorithm originally used by Mersch et al. (2013; for the task of functional group allocation). Community membership assignments, compared at an individual node level for a given network, resulted in an average 90.13 ± 7.25% similarity between the two methods across all the networks in the dataset. While the functional group assignments used in our analyses were taken directly from Mersch et al.’s analysis (which were validated by behavioural observations), this similarity of community assignment is important as we wanted to certify that the structure of communities detected by both algorithms was not divergent. The bridge betweenness metric used the communities from the Louvain algorithm as it provided a substantial reduction in computational time, and was indicative of the potential for an ant to connect ants from different functional groups because of the high similarity with the community assignments used to determine functional groups (i.e. by infomap and Mersch et al., 2013).

### Individual network centrality measures and task switching

We compared network measures and their variances across each of the functional groups for three categories: overall across all ants, for just switching ants, and for just ants that remained consistent in their tasks during the assessment periods (summarised in Figure 2; all values listed in Supplementary material 2 Table S2 and additional visualisation in Figure S2), relative to their respective reference networks. Variance was assessed due to apparent substantial fluctuations in the metrics for ants that switched within the 10-day period leading to a task assessment point (Figure S3). These fluctuations are represented by the 95% confidence intervals of the variances which indicate how stable the relative network measures were across colonies and over time (i.e., a larger confidence interval represents more fluctuations and less stability).

**Figure 2:**
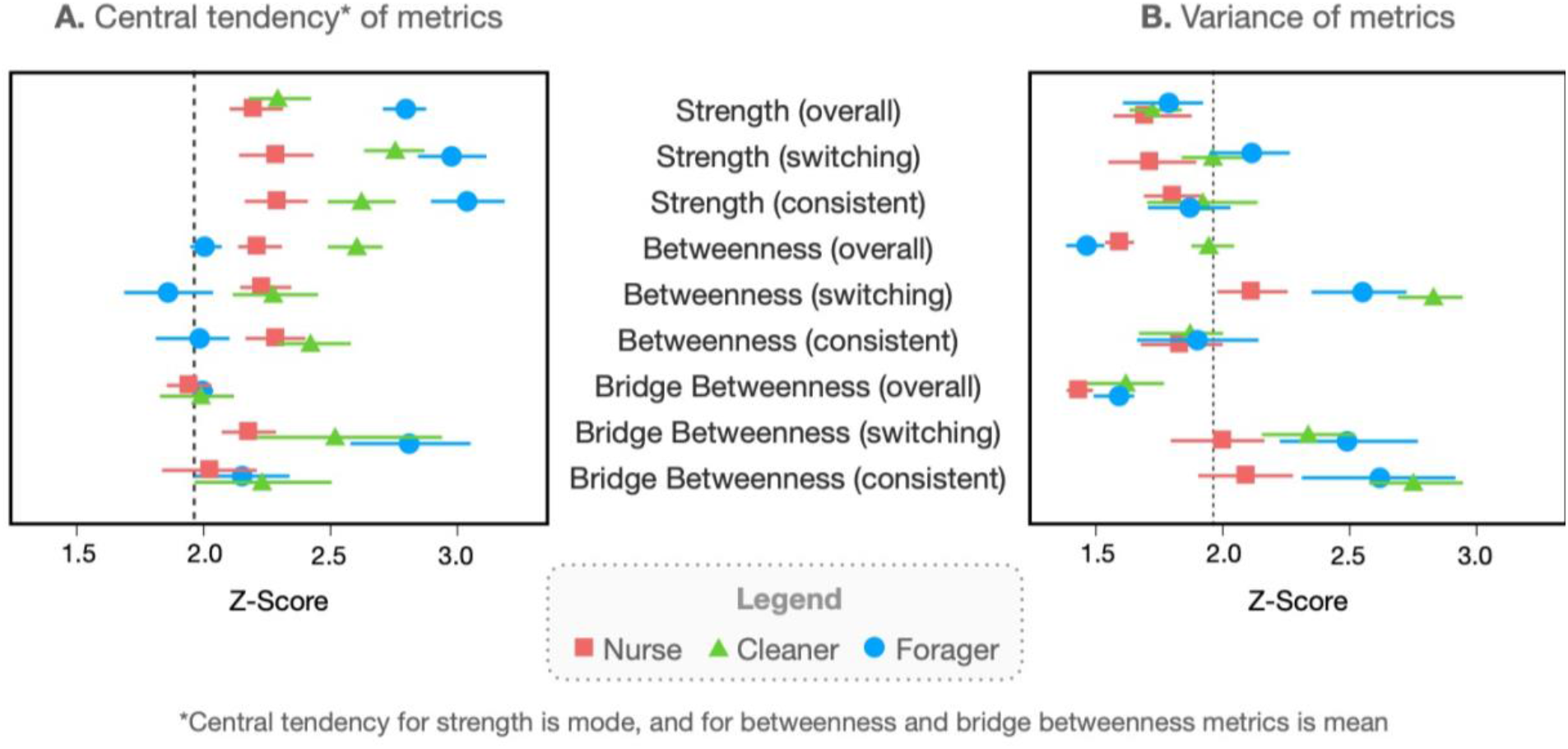
Z transformed values of the central tendencies (A) (note that the central tendency for strength is mode while for the other two metrics is mean) and variances (B) of the strength mode, betweenness and bridge betweenness determined for all ants across the six colonies. For each functional group, values were determined for all ants (overall), just ants that changed functional group during assessment periods (switching), and those ants that performed the same tasks from one time period to the next (consistent). Values were determined for every observed ant network (each colony, for each day) and for each of the reference networks before being Z-transformed to facilitate comparisons. Points are the values for each functional group (points depict average value and bars represent 95% confidence intervals among the measured central tendencies and variances (network-level statistics) of observed networks across days and colonies): nurses (red squares), cleaners (green triangles) and foragers (blue circles). The dashed black lines represent the upper extent of 95% confidence interval of the same normalised metrics from reference network simulations. (See Figure S2 for a more complete representation).

When we compared the distribution of the strength metric using the mode across each of the tasks, we found that foragers had the highest strength mode of any of the groups across all three of the categories, showing that they had the most frequent interactions over a day regardless of whether they remained foragers or switched task at some point. Values of variance (relative to their respective reference networks) of the overall strength did not significantly differ across functional groups or from the reference networks (see the 95% confidence intervals in Figure 2B) and the mode of strength remained fairly consistent across functional groups (see the 95% confidence intervals in Figure 2A). When we looked at strength just for switching ants, we found that the mode differed significantly across functional groups and was significantly greater than for the reference networks. Out of these, foragers that switched had the highest strength mode. Strength mode variance of switching ants did not vary significantly among functional groups. However, variance of the strength mode of switching foragers was higher and more variable (i.e. larger confidence interval) than the reference networks, indicating less stability of this metric among these individuals and over time. When we looked at strength just for ants that were consistent, we found that the mode and variance followed the same pattern seen for ants that switched, i.e., consistent foragers had higher strengths and the values for all groups were of the same magnitude as those for the ants that switched.

At the colony level, the betweenness metric was stable (i.e., confidence intervals were small for the both mean and the variance relative to the reference networks) and cleaners played the most important role in connecting individual ants for flow of information, as they had significantly higher betweenness than nurses and foragers (Figure 2). When we assessed betweenness just for ants that switched, we found that mean betweenness centrality measures were significantly greater than those for the reference networks, except for foragers. Betweenness of switching ants was more variable than for consistent ants. Consistent ants had the same relative patterns and magnitude of mean betweenness centrality as ants that switched: consistent cleaners and nurses had higher mean betweenness than consistent foragers. However, the variance of betweenness was no longer significantly different than the reference networks, thus consistent ants maintained a less variable betweenness distribution among networks and through time (i.e., the mean and variance remain within small confidence intervals, and that the variance is not significantly different from reference networks) than switching ants (where even if the mean has similar values, the variance is much higher than reference networks).

Since the communities we detected mapped primarily onto the previously determined functional groups (see results above), a high bridge betweenness indicated a high potential for connecting functional groups in a colony. When we compared bridge betweenness across each of the functional groups at the colony level, we found that the overall mean bridge betweenness values and their variance did not vary among the functional groups or from the reference networks, indicating that connections among network group communities stayed stable among networks and through time at the colony level. Mean bridge betweenness was higher for the switching ants than for the consistent ants for all functional groups, though only significantly higher for foragers. All ants that switched had significantly higher mean bridge betweenness than the overall colony values per functional group, suggesting that ants that switched played an important role in connecting communities for information flow in the colony. The mean bridge betweenness of consistent ants did not vary significantly among the functional groups or from the reference network distribution. Although the variance of cleaners and foragers for both ants that switched and those that were consistent was significantly higher and more variable than the reference networks, the overall colony variance values remained stable with small confidence intervals; these results may indicate that these interaction structures could be important for colony cohesion at the community level.

### Global information flow and task switching

We measured Effectiveness, the normalised Effective Information (difference between how deterministic and degenerate a network is), as a function of the proportion of nurses, cleaners or foragers in each colony for each day (resulting in 246 Effectiveness measures, Figure 3). We found that the colony networks with high proportions of nurses and cleaners had higher Effectiveness, but that the dependencies based on the linear model were weak and non-significant (adj. R^2^=0.12, P = 0.063, Figure 3A for nurses; and adj. R^2^=0.11, P = 0.052, Figure 3B for cleaners). Effectiveness significantly decreased with increasing proportions of foragers in a colony (adj. R^2^=0.22, P = 0.037, Figure 3C). This negative relationship between the proportion of foragers and colony-level Effectiveness suggests that interactions involving foragers were noisier than those involving only nurses or cleaners.

**Figure 3:**
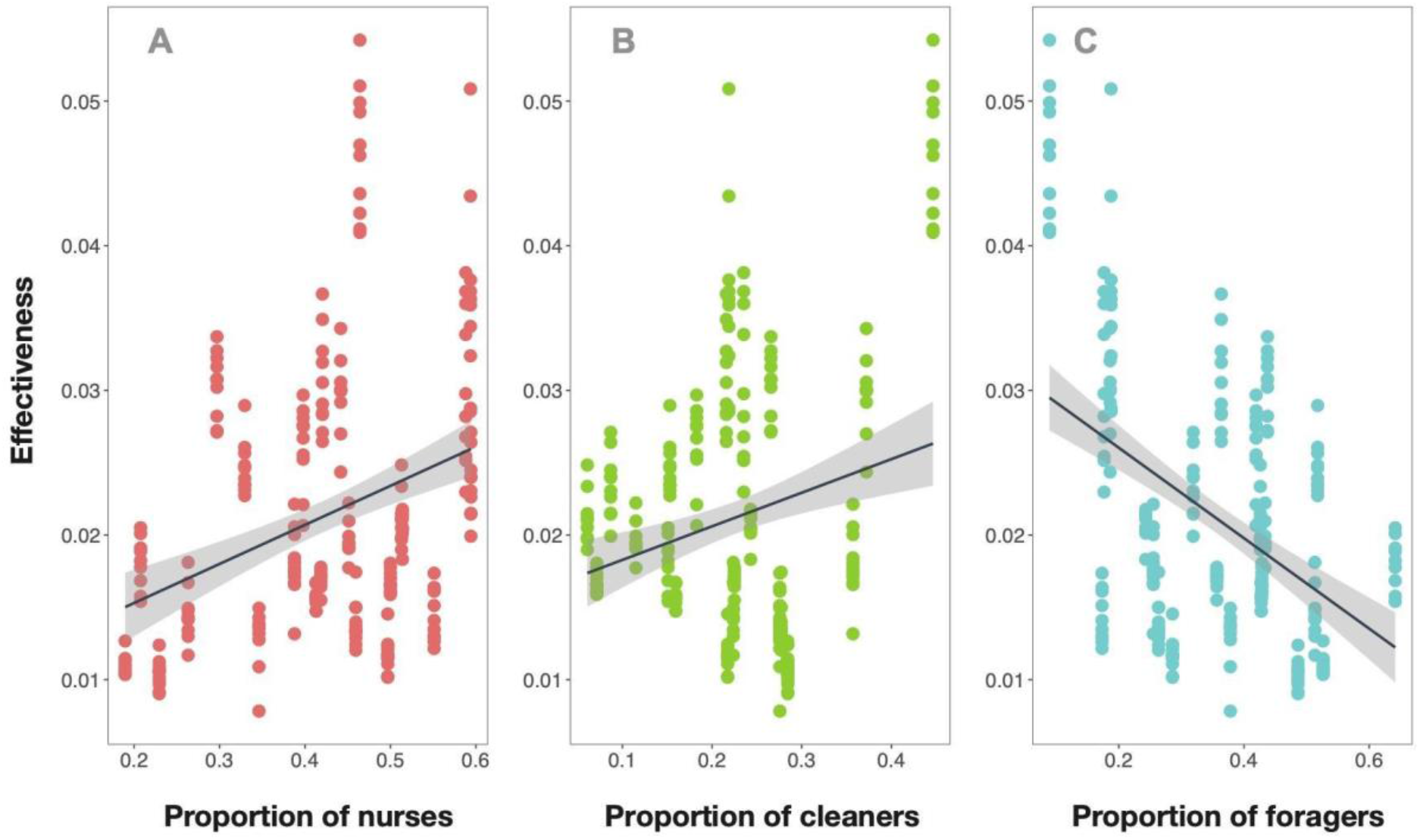
Effectiveness (normalised Effective Information) of the interaction networks constructed for each colony and every day of the experiment as a function of the proportion of different functional groups in the networks. Data are stacked because the available granularity for task allocation was at a 10-day interval. Linear models fit to Effectiveness as a function of the proportions of nurses (A) and cleaners (B) separately return a nominally positive dependence (adj. R^2^=0.12, p = 0.063 for nurses; and adj. R^2^=0.11, p = 0.052 for cleaners). Effectiveness as a function of the proportion of foragers (C) returns a strong negative dependence (R^2^ =0.22, p = 0.037).

### Task interaction matrix and task switching

We tested whether previous interaction patterns affected switching behaviour using a task interaction matrix. We found that ants that remained consistent in their tasks usually interacted most with other ants occupying their same task (Table 1, Consistent ants). For example, consistent nurses were significantly more likely to only have interacted with other nurses (90% of nurse interactions, p=0.0326). Although cleaners and foragers who stayed within their functional group also more commonly interacted with other cleaners or foragers, this difference in interaction frequency was not significantly higher than expected by chance. For simplicity and comparison throughout networks, we present the average of proportion of interactions in each case in Table 1.

**Table 1:**
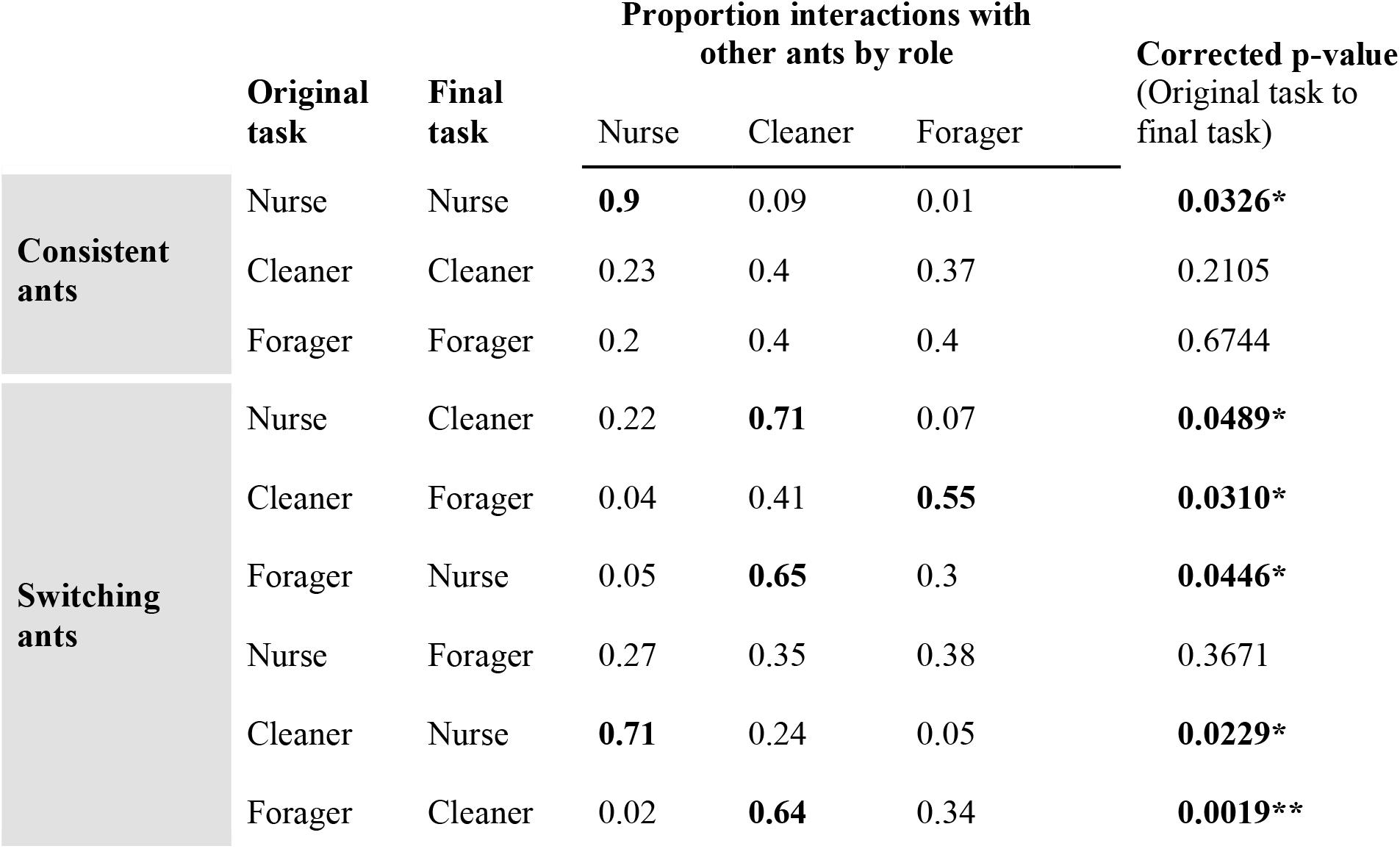
The task interaction matrix, showing the proportion of an ant’s interactions with a specified functional group before switching from its original to final group. P-values were calculated using a chi-square test contrasting the observed interaction proportions with the reference model results for each type of task transition; values significantly different from those obtained from their reference networks (after multiple comparison corrections) are indicated with asterisks. Bold type indicates the task and proportion of interactions with ants of that task that were dominant in each category (and which were higher than expected by random chance).

However, most ants that switched to a new task interacted with ants occupying a different task prior to switching (Table 1, Switching ants). For example, nurses who switched to cleaning had interacted more frequently with cleaners (71% of nurse interactions) and this was significantly more likely to occur based on interaction history than by random chance (p = 0.0489). The result that an ant would transition to a group that it previously interacted with the most was significant for the following other transitions: cleaner to nurse, cleaner to forager, and forager to cleaner. Interestingly, foragers who switched to nursing were significantly more likely to have interacted more with ants of a different functional group, the cleaners (65% of forager interactions who then switched to nursing). However, it is important to note that this forager to nurse transition only occurred in a few cases in the experimental data, so these results should be interpreted with caution (due to the low number of observed ants), even though the interaction frequency data was sufficient for the statistical comparison (see supplementary files for sample size information).

## DISCUSSION

We explored task allocation in ant colonies to determine whether we could explain how ants switched tasks based on information flow among functional groups and the interaction history of the individuals. Mersch et al. (2013) determined that task switching was a noisy process with a lot of individual variation, but that at least some of the task switching could be explained by age polyethism based on the spatial division of workers mediating the structure of the interaction network. In our analyses, we focused specifically on this noisy process of task switching. Our approach allowed us to determine that previous interaction history can help explain some of the noise behind task switching in *Camponotus fellah* colonies and provides novel insight into task switching behaviour in these ants.

Our results suggest that ants in different functional groups had varying levels of importance for information flow between individuals and groups in a colony, based on their individual roles in network connectivity as determined by the node-level metrics. Ants that switched tasks often occupied positions in the interaction network that had high potential for supporting information flow between groups. Network analyses, combined with the task interaction matrix, allowed us to describe how the architecture of interactions was related with the distribution of and switching among tasks in an ant colony.

At the scale of ant-to-ant interactions, we found that ants classified into the three main tasks (forager, cleaner, nurse) differed in how they interacted with each other, which affected their roles in information flow for the colony. Foragers had the highest interaction strength mode – they interacted more frequently than cleaners and nurses. Cleaners, however, had higher betweenness and thus were key connections between ants interacting in the colony.

Ants that switched tasks functioned as key connectors for information flow in the colony, supporting colony cohesion. In general, mean betweenness was higher for ants that switched than for ants that remained consistent in their task, although confidence intervals overlapped. Bridge betweenness (which indicated how ants connected different communities within the colony) was significantly higher for ants that switched. This suggests that ants who switched tasks throughout the course of the experiment, and particularly foragers, played an important role in connecting functional groups through information flows. Their high bridge betweenness means that they occupied a key network position for receiving and transmitting information before they switched tasks. If learning is required when ants switch tasks, this increased access to information may have allowed them to learn new behaviours more quickly, helping them transition to a new task. In general, cleaners were less likely to interact within their functional group (which was consistent with Mersch et al.’s 2013 results). The low group cohesion of cleaners may strengthen colony-wide cohesion.

The variability of the centrality metrics may be related to cleaners’ and foragers’ ability to transition tasks. Cleaners and foragers who switched functional groups had significantly higher variances of betweenness and bridge betweenness within networks. Across networks and over time, these variances also had a larger range, showing that these individual measures of social network connectivity changed more and were overall less consistent among individuals. However, when all ants in a colony were grouped for calculating the node-level network metrics, pooled variances were not higher than those for the reference networks and had small confidence intervals. So, while these metrics varied significantly among functional groups and when ants switched tasks, overall information flow in a colony remained fairly stable and colony cohesion was maintained.

At the group level, the operationalisation of Effective Information as a measure of the noisiness of network mechanisms is relatively new and under-explored. Our results for the six *Camponotus fellah* colonies show a correlation between variations in Effectiveness and the functional group composition of each colony. We found that a higher proportion of foragers led to noisier potential communication among ants. Paired with the results on interaction strength, this means that foragers interacted more frequently than ants performing other tasks and that they had more diverse interactions with ants at different positions in the interaction network. Results on centrality measures and Effectiveness can be linked with task allocation through our task interaction matrix. The matrix shows how previous interactions with ants in a given task are associated with a higher probability of the ant switching to that task. These results are consistent with previous work in another species: Gordon and Mehdiabadi (1999) found that, in red harvester ants, ants switching from other tasks to midden work were more likely to have interacted with midden workers, and that switching was more likely to occur the more frequent those interactions. In our results, interactions with foragers were correlated with switches to foraging: both cleaners and nurses who switched to foraging had a higher probability of interacting with foragers. Switches from foragers to other tasks, however, showed different dynamics. Both foragers who switched to nursing and foragers who switched to cleaning had a higher probability of interacting with cleaners. These results should be interpreted with some caution because ants switching from foragers to nursing was only observed three times (the interaction frequency data, however, was sufficient for the chi-square comparison). Consistent with betweenness results, these switching results show that cleaners were central in driving switching patterns by occupying key positions for information flow in the networks. These patterns suggest that, while previous interaction patterns were correlated to switching behaviour, the degree of correlation varied depending on the role played by the interacting ants and on the overall information flow of the system. It is important to note that without more detailed data we cannot determine whether a change in task or change in interactions happened first, but these insights provide valuable information about system dynamics and suggestions for future experiments.

In future research, it would be interesting to further explore task switching in systems with a higher granularity of data collection across both behaviours and interactions. One limitation to the current analysis is that the task each ant was assigned to is assessed based on the interaction patterns, not the types of actions or tasks the ant completes in the colony. Even though interaction community membership was paired with behavioural observations by Mersch et al. (2013), it may not have been at the level of detail needed to assess fine-grain interaction patterns and task performance. It would be interesting to use the combination of network methods and behavioural observations to further explore existing results on the relationship between repetition (Langridge et al., 2004) and the existence of experienced individuals (Langridge et al., 2007) on task performance. Assessing not just who an ant interacts with, but what actions that ant is actually completing, would provide useful additional insight into the timing of behavioural and social change. This kind of data would allow researchers to determine whether an ant alters its behaviours first (for example, decreasing cleaning behaviours and increasing nursing behaviours) which then results in a change in the social interaction patterns, or whether an ant first begins to change its social interaction patterns (for example, interacting less with other cleaners and more with nurses) and then alters its behaviour from cleaning actions to nursing actions. Future targeted data collection, involving both social and behavioural observations, paired with statistically robust network methods, could be used to further explore the relationships between patterns of interactions, individual-level behaviour, and group-level behaviour.

## ACKNOWLEDGEMENTS

We sincerely thank the organizers of 2019 Complex Networks Winter Workshop (CNWW) for bringing us together and Brennan Klein for helpful discussions. A.S.’s contribution to this research was supported in part through training by NSF award DGE-1632976. S.W.’s contribution was supported in part by an NSF-GRF. EAH was supported by NSF IOS 2015932 during preparation of this work.

## Author contributions

All the authors conceived the idea and the methodology together; AS and SW did the analysis, and all the authors wrote the manuscript.

## Data availability

All scripts and data used in this project are available at https://github.com/anshuman21111/ant-colony-networks. A reformatted version of the Mersch et al. (2013) dataset can be found as Supplementary material 1.

