## Supplementary figures and tables for "Interactions and information: Exploring task allocation in ant colonies using network analysis"

**Supplementary Material 2**

**Table S1**: Overview of colony composition.

| **Colony** | **Group** | ***t1*** | ***t4*** | **Colony** | **Group** | ***t1*** | ***t4*** |
| --- | --- | --- | --- | --- | --- | --- | --- |
| Colony 1 | N | 25 | 11 | Colony 4 | N | 38 | 19 |
|  | C | 32 | 8 |  | C | 35 | 7 |
|  | F | 52 | 34 |  | F | 25 | 6 |
|  | Q | 1 | 1 |  | Q | 1 | 1 |
|  | NA | 0 | 56 |  | NA | 0 | 66 |
|  | total | 110 | 110 |  | total | 99 | 99 |
| Colony 2 | N | 70 | 26 | Colony 5 | N | 68 | 19 |
|  | C | 35 | 25 |  | C | 41 | 17 |
|  | F | 22 | 5 |  | F | 39 | 28 |
|  | Q | 1 | 1 |  | Q | 1 | 1 |
|  | NA | 0 | 71 |  | NA | 0 | 84 |
|  | total | 128 | 128 |  | total | 149 | 149 |
| Colony 3 | N | 38 | 37 | Colony 6 | N | 80 | 37 |
|  | C | 93 | 17 |  | C | 35 | 19 |
|  | F | 25 | 39 |  | F | 46 | 32 |
|  | Q | 1 | 1 |  | Q | 1 | 1 |
|  | NA | 0 | 63 |  | NA | 0 | 73 |
|  | total | 157 | 157 |  | total | 162 | 162 |

**Table S2:** Network metrics (mean and variance) for each functional group^†^.

| **Property** | **Type** | **Nurses** | **Cleaners** | **Foragers** |
| --- | --- | --- | --- | --- |
| Strength | Switching mean | 2.17 ≤ **2.35** ≤ 2.52 | 2.76 ≤ **2.92** ≤ 3.05 | 3.01 ≤ **3.17** ≤ 3.33 |
|  | Switching variance | 1.46 ≤ **1.65** ≤ 1.88 | 1.8 ≤ **1.96** ≤ 2.1 | 1.97 ≤ **2.14** ≤ 2.32 |
|  | Consistent mean | 2.21 ≤ **2.35** ≤ 2.48 | 2.59 ≤ **2.74** ≤ 2.91 | 3.08 ≤ **3.25** ≤ 3.43 |
|  | Consistent variance | 1.63 ≤ **1.76** ≤ 1.92 | 1.66 ≤ **1.85** ≤ 2.16 | 1.65 ≤ **1.92** ≤ 2.04 |
|  | Overall mean | 2.12 ≤ **2.25** ≤ 2.38 | 2.21 ≤ **2.35** ≤ 2.5 | 2.84 ≤ **2.96** ≤ 3.06 |
|  | Overall variance | 1.49 ≤ **1.66** ≤ 1.85 | 1.54 ≤ 1.68 ≤ 1.8 | 1.56 ≤ **1.75** ≤ 1.92 |
| Betweenness | Switching mean | 2.17 ≤ **2.29** ≤ 2.43 | 2.14 ≤ 2.33 ≤ 2.53 | 1.63 ≤ **1.83** ≤ 2.05 |
|  | Switching variance | 1.98 ≤ **2.14** ≤ 2.32 | 2.84 ≤ **3.01** ≤ 3.14 | 2.43 ≤ **2.67** ≤ 2.88 |
|  | Consistent mean | 2.19 ≤ **2.33** ≤ 2.46 | 2.37 ≤ **2.5** ≤ 2.7 | 1.76 ≤ **1.97** ≤ 2.12 |
|  | Consistent variance | 1.64 ≤ **1.81** ≤ 2.03 | 1.81 ≤ **1.87** ≤ 2.02 | 1.62 ≤ **1.88** ≤ 2.19 |
|  | Overall mean | 2.15 ≤ **2.25** ≤ 2.37 | 2.58 ≤ **2.73** ≤ 2.84 | 1.95 ≤ **2.01** ≤ 2.08 |
|  | Overall variance | 1.45 ≤ **1.51** ≤ 1.57 | 1.85 ≤ **1.94** ≤ 2.06 | 1.25 ≤ **1.36** ≤ 1.44 |
| Bridge betweenness | Switching mean | 2.09 ≤ **2.22** ≤ 2.35 | 2.27 ≤ **2.63** ≤ 3.15 | 2.69 ≤ **2.97** ≤ 3.25 |
|  | Switching variance | 1.76 ≤ **1.99** ≤ 2.21 | 2.19 ≤ **2.41** ≤ 2.64 | 2.28 ≤ **2.59** ≤ 2.92 |
|  | Consistent mean | 1.8 ≤ **2.03** ≤ 2.25 | 1.95 ≤ **2.27** ≤ 2.59 | 1.96 ≤ **2.18** ≤ 2.41 |
|  | Consistent variance | 1.89 ≤ **2.11** ≤ 2.34 | 2.68 ≤ **2.91** ≤ 3.13 | 2.38 ≤ **2.75** ≤ 3.09 |
|  | Overall mean | 1.82 ≤ **1.93** ≤ 2.04 | 1.79 ≤ **1.98** ≤ 2.14 | 1.94 ≤ **1.99** ≤ 2.05 |
|  | Overall variance | 1.26 ≤ **1.32** ≤ 1.38 | 1.36 ≤ **1.55** ≤ 1.73 | 1.41 ≤ **1.51** ≤ 1.59 |

### ^†^Z-transformed values of various functional groups for all the six colonies and all days. Mean values are in bold and shown within the range of their 95% confidence intervals.

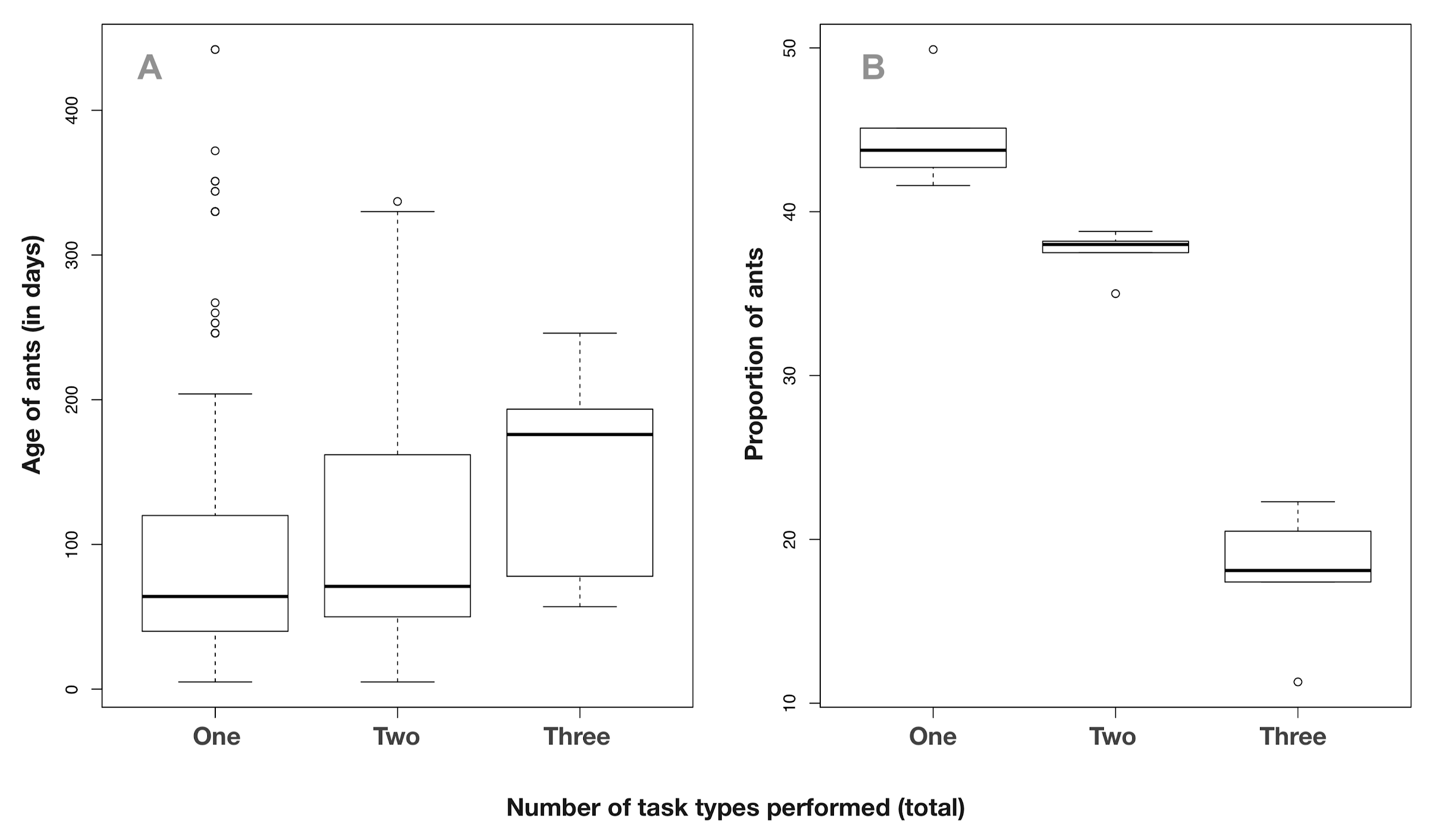

**Figure S1:** Age polyethism (A) and switching cost (B). We checked the hypothesis that ants become more flexible with age, by plotting the number of switches	against the age of the ant. The plot does not show	statistically significant results, but it shows	that there may be a correlation between ant age and switching frequency (with a p value of 0.23 between one-two, 0.27 between two-three and 0.08 between one-three), which would require more data points to be confirmed. In S1B, we show the likelihood that an ant will stay in the same task throughout the experimental time versus performing two or three tasks (with the possibility of switching between tasks). Ants are more likely to stay within a single task, confirming the existence of a switching cost.

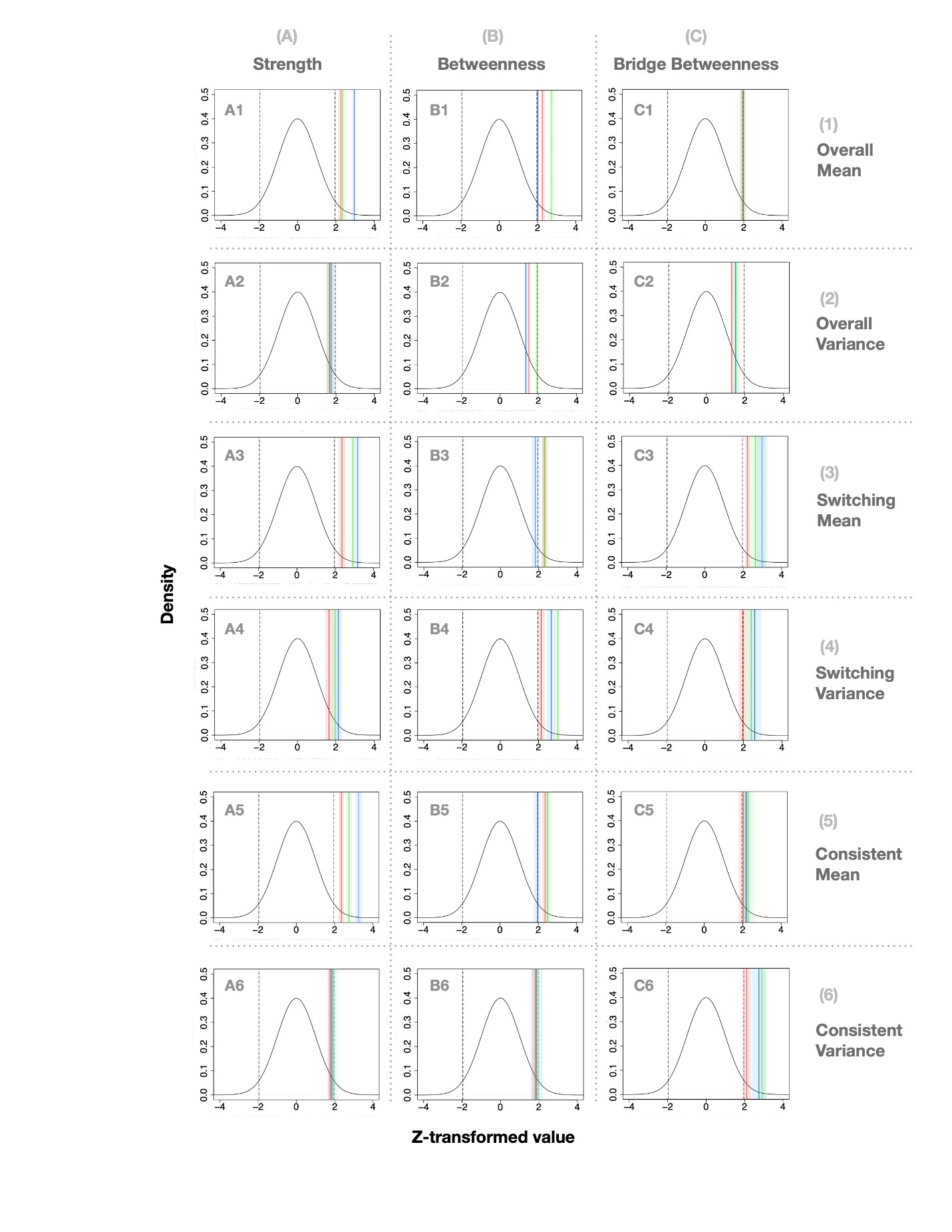

**Figure S2:** Z transformed values of the mean and variances of the strength, betweenness and bridge betweenness (columns A, B and C respectively) determined for all ants across the six colonies. The first two rows (rows 1 and 2; A1-C2) are values for all the ants in the experiment (overall colony values). The next two rows (rows 3 and 4; A3-C4) show values for ants that switched during the experiment, and the last two rows (rows 5 and 6; A5-C6) show values for for consistent ants (no switching) during the experiment. Line colours correspond to ant functional group type: nurses (red), cleaners (blue), and foragers (green). Shaded areas are the 95% confidence intervals based on seeded random networks. The null distribution created from the null network simulations is shown in black with a marked confidence interval (dashed lines).

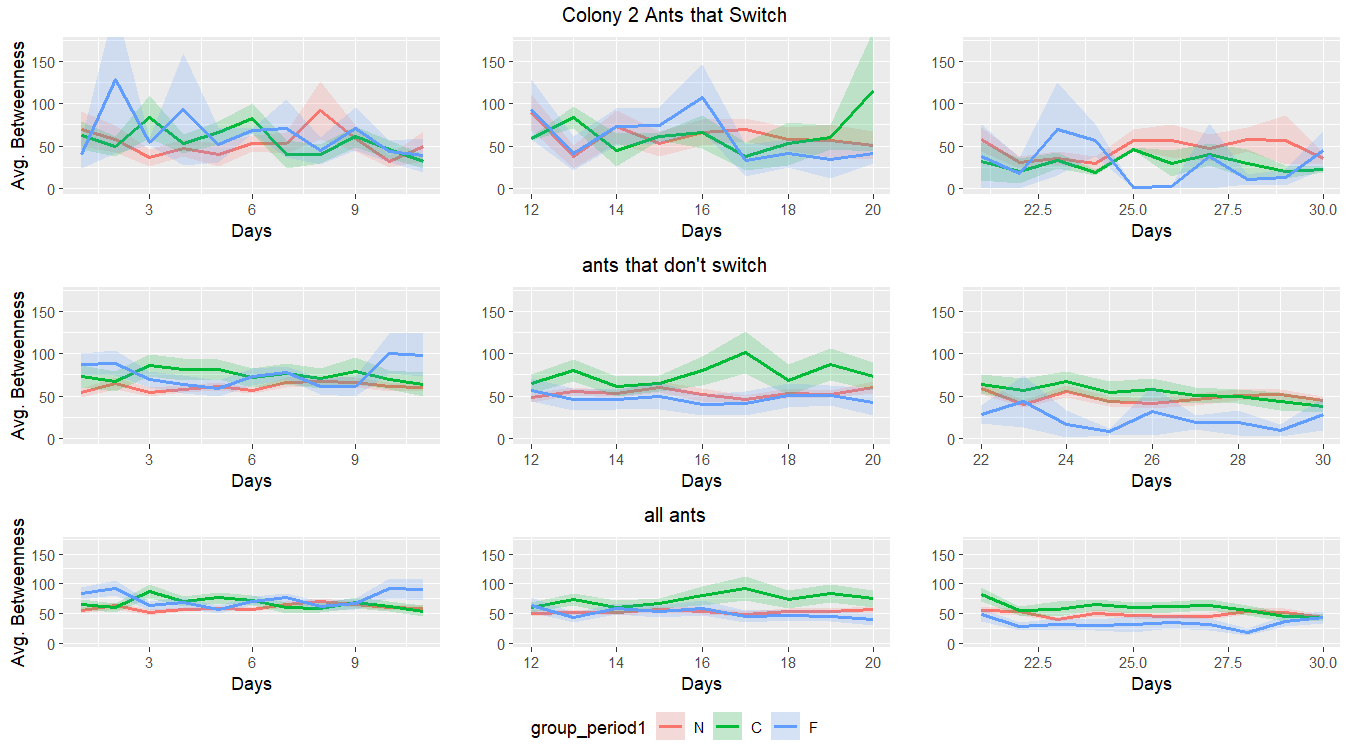
**Figure S3:** Daily values of average node betweenness (shaded area represents standard deviation) grouped for nurses (red), cleaners (green), and foragers (blue) in Colony 2, split by ants that switched (top row), ants that stayed consistent (middle row), and all ants (bottom row). Each plot is a 10 day period leading up to a task assessment point. Average betweenness during the 10 day periods varied more for ants that switched, prompting inclusion of variance in the main analyses. Similar results were observed for the other colonies and metrics (strength and bridge betweenness) and additional figures are available on the GitHub repository for the paper.
